## Supplementary text for "Live Cell Imaging of Single RNA Molecules with Fluorogenic Mango II Arrays"

**Supplementary Table 1.**

| Construct | Sequence (5' → 3') |
|---|---|
| Mango II | GGCACGTACG AAGGAGAGGA GAGGAAGAGG AGAGTACGTG C |
| M2 x3 array (5 nt) | GGCACGTACG AAGGAGAGGA GAGGAAGAGG AGAGTACGTG CTTTTTGGAC ATGCCGAAGG AGAGGAGAGG AAGAGGAGAG GCATGTCCTT TTTGGGTGCA TCGAAGGAGA GGAGAGGAAG AGGAGAGATG CACCC |
| M2 x3 array (1 nt) | GGCACGTACG AAGGAGAGGA GAGGAAGAGG AGAGTACGTG CTGGACATGC CGAAGGAGAG GAGAGGAAGA GGAGAGGCAT GTCCTGGGTG CATCGAAGGA GAGGAGAGGA AGAGGAGAGA TGCACCC |
| Broccoli | GGAGACGGTC GGGTCCAGAT ATTCGTATCT GTCGAGTAGA GTGTGGGCTC C |
| Broccoli Double (10 nt) | GGAGACGGTC GGGTCCAGAT ATTCGTATCT GTCGAGTAGA GTGTGGGCTC CCTCTCTCTC TGGAGACGGT CGGGTCCCTC GATTCGTCGA GGTCGAGTAG AGTGTGGGCT CC |
| Broccoli Triple (5 nt) | GGAGACGGTC GGGTCCAGAT ATTCGTATCT GTCGAGTAGA GTGTGGGCTC CTTTTTGGCT ACGGTCGGGT CCCTCGATTC GTCGAGGTCG AGTAGAGTGT GGGAGCCTTT TTGGTCACGG TCGGGTCCTC TAATTCGTTA GAGTCGAGTA GAGTGTGGGG ACC |
| M2-AF488 smFISH_1 | AF488N//CGGCATGTCC AAAAAGCACG |
| M2-AF488 smFISH_2 | AF488N//ATGCACCCAA AAAGGACTG |
| M2-AF488 smFISH_3 | AF488N//TACGTGCCAA AAAGGGTGCA |
| MS2v5-FAM smFISH_1 | FAM//TGATTGTGAAGTGTCGGGTG |
| MS2v5-FAM smFISH_2 | FAM//TCCACCCTTGTGTATTGTAC |
| MS2v5-FAM smFISH_3 | FAM//TGTAATGTGTCTGGAGGGTG |
| MS2v5-FAM smFISH_4 | FAM//GCTTCTGTTTGATTGGATTT |
| MS2v5-FAM smFISH_5 | FAM//GATGGTGATTCCTTGTTGTA |
| MS2v5-FAM smFISH_6 | FAM//GTATATTGCACAGGGAATCC |
| MS2v5-FAM smFISH_7 | FAM//GATATTCGGGAGGCGTGATC |
| MS2v5-FAM smFISH_8 | FAM//ACGCACTGAATTCGAAAGCC |

| | |
|---|---|
| MS2v5-FAM smFISH_9 | FAM//ATTCGACTCTGATTGGCTGC |
| MS2v5-FAM smFISH_10 | FAM//CTCTTCGCGAAAGTCGACTT |
| MS2v5-FAM smFISH_11 | FAM//TAAGAATGGCGCGAAGGCTG |
| MS2v5-FAM smFISH_12 | FAM//GTAGGGGAGAGTGTGGTTTG |
| MS2v5-FAM smFISH_13 | FAM//CAGGAACGCTGATGCTGTTC |
| MS2v5-FAM smFISH_14 | FAM//TTTTCTTGAGTTGGGTACTG |
| MS2v5-FAM smFISH_15 | FAM//TGATGCTGCATGGGGACATA |
| MS2v5-FAM smFISH_16 | FAM//TTGGGGATGTATTCTTGGGG |
| MS2v5-FAM smFISH_17 | FAM//TTGGTGCTCGGATGTGATTT |
| MS2v5-FAM smFISH_18 | FAM//AAGAAACAACACTCCGAGCC |
| MS2v5-FAM smFISH_19 | FAM//ATGGAGGGTTTGTCCAGTTG |
| MS2v5-FAM smFISH_20 | FAM//TTTGTCTTGTTGGTGAGAGT |
| MS2v5-FAM smFISH_21 | FAM//CTGATGCTGCTTCGAGAAGA |
| MS2v5-FAM smFISH_22 | FAM//GTATGCTCGAGTGTTTCGAA |
| MS2v5-FAM smFISH_23 | FAM//GATCGTCCACCCAAGAAATA |
| MS2v5-FAM smFISH_24 | FAM//AATTCGTGAGAGCATGGGTG |
| MS2v5-FAM smFISH_25 | FAM//TCGTATTGGACGTGGAACGA |
| MS2v5-FAM smFISH_26 | FAM//TCGTGATCCCGAAAGGTAAG |
| MS2v5-FAM smFISH_27 | FAM//ATCGTGCATGCTTGAATGTC |
| MS2v5-FAM smFISH_28 | FAM//GTTGAGACTTGTGGAGCATG |

| | |
|---|---|
| MS2v5-FAM smFISH_29 | FAM//TGAACCCATTTGGTAGTTTC |
| MS2v5-FAM smFISH_30 | FAM//TTTGAGGTAGGAGTGGGTTC |
| MS2v5-FAM smFISH_31 | FAM//TTGCCAGTTTTGTGGGAAGA |
| MS2v5-FAM smFISH_32 | FAM//TTTGGTATGTTGGAATGGGC |
| MS2v5-FAM smFISH_33 | FAM//GATGCTGTACCAGTAATTGT |
| MS2v5-FAM smFISH_34 | FAM//TAGTAGTGAGAGATGTGGGC |
| MS2v5-FAM smFISH_35 | FAM//TGCTGAACGGTTTGGTTTTT |
| MS2v5-FAM smFISH_36 | FAM//TTGATTTTTCCGTGTGTACC |
| MS2v5-FAM smFISH_37 | FAM//GTCTTTCGTATTTGTAAACC |
| MS2v5-FAM smFISH_38 | FAM//TTGCGCTGGACGAAAGCGTG |
| MS2v5-FAM smFISH_39 | FAM//CCGTCGGATGTTTTTCGTAA |
| MS2v5-FAM smFISH_40 | FAM//GGTTGTAAGTTTGTGGGTTG |
| MS2v5-FAM smFISH_41 | FAM//TTGATGTACGGTGCGGTGAT |
| MS2v5-FAM smFISH_42 | FAM//ATCGATATGAGATCTGAGGT |

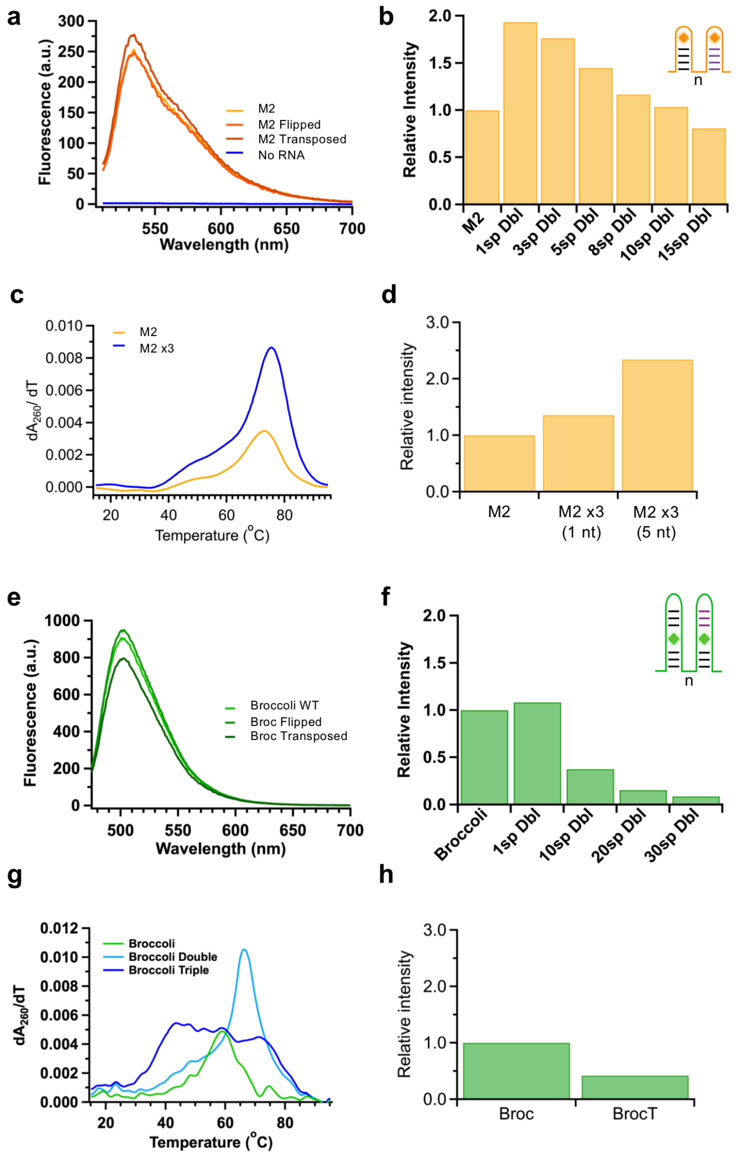


**Fig. S1| *In vitro* characterisation of aptamer arrays**

(**a**) Fluorescent spectra of individual mutant Mango sequences (40 nM) either flipped or transposed in the presence of TO1-Biotin (200 nM). (**b**) Relative fluorescent intensity of dimeric Mango constructs with alteration of the nucleotide spacing (n) between adjacent aptamers. (**c**) Differentials of the UV melting curves for Mango II and the Mango II x3 (5nt) array. Adapted from main figure 1 to aid comparison with Broccoli constructs. (**d**) Relative fluorescent intensity of Mango II and the Mango II x3 (1 nt) and (5nt) arrays. (**e**) Fluorescent spectra of individual mutant Broccoli sequences (40 nM) with the P3 stem either flipped or transposed in the presence of DFHBI (1 μM). (**f**) Relative fluorescent intensity of dimeric Broccoli constructs with alteration of the nucleotide spacing (n) between adjacent aptamers. (**g**) Differentials of the UV melting curves for Broccoli the Broccoli (10 nt) dimer and the Broccoli trimeric array. (**h**) Relative fluorescent intensity of Broccoli and the Broccoli trimeric array (BrocT).

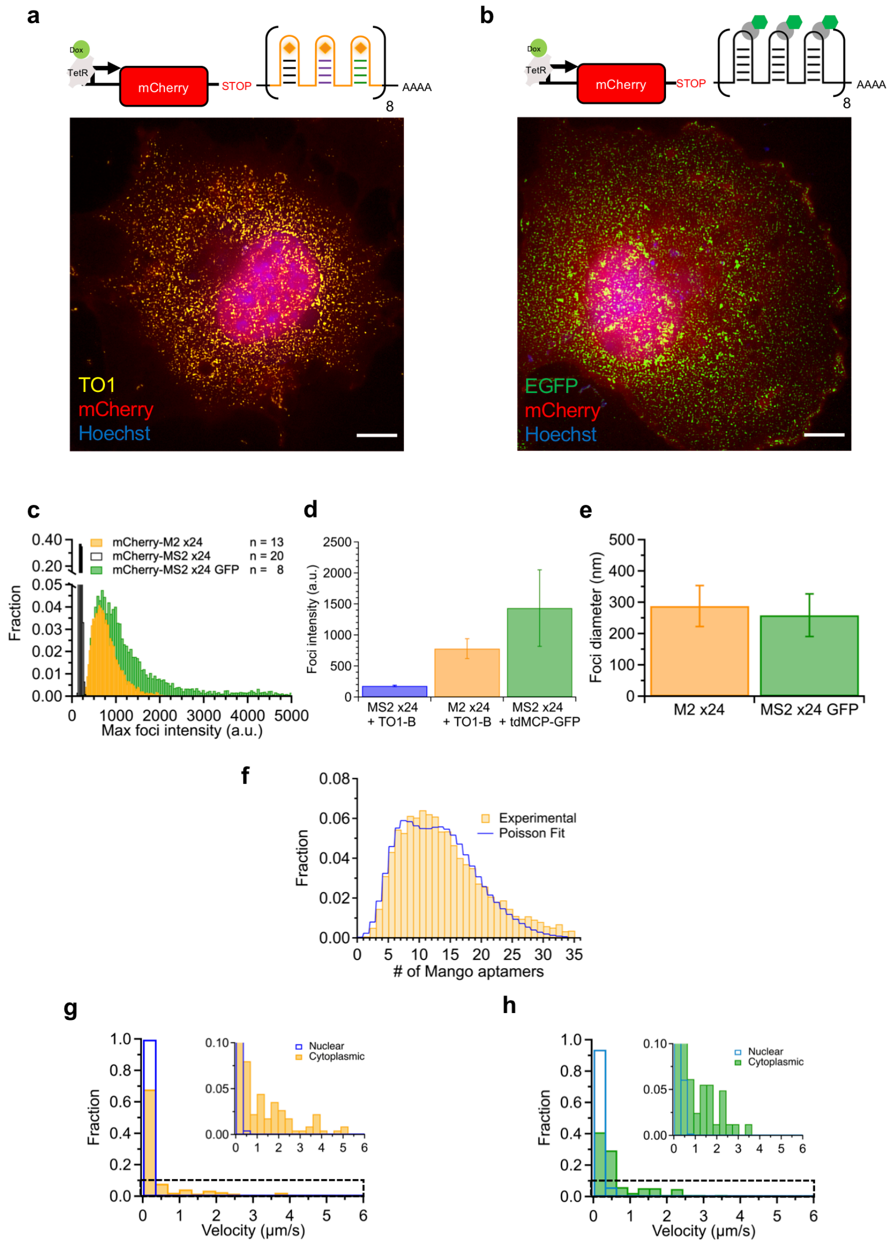


**Fig. S2 | Comparison of M2 x24 and MS2v5 x24 labelled single RNA molecules**

(**a**) Schematic of the Dox inducible mCherry-M2 x24 construct and a representative maximum projection of a cell displaying evenly dispersed single molecules. Cells stained with 200 nM TO1-B. (**b**) Schematic of the Dox inducible mCherry-MS2v5x24-tdMCP-EGFP construct and a representative maximum projection of a cell displaying evenly dispersed single molecules. Both scale bars = 10 μm, in Cos-7 Cells. (**c**) Distribution of single molecule intensities for cells expressing the mCherry-M2 x24 construct + TO1-B (200 nM - yellow), mCherry-MS2v5 x24 + TO1-B (200 nM - black) and mCherry-MS2v5 x24-tdMCP-EGFP (green), 7008, 30,429 and 10,338 foci respectively. N shown in figure depicts number of cells used for analysis. Figure adapted from main figure 2f for direct comparison with EGFP foci. (**d**) Mean foci intensity of each construct with the error shown as standard deviation. (**e**) Mean foci diameter of M2 x24 and MS2v5 x24-tdMCP-EGFP RNAs with the error shown as standard deviation. (**f**). Foci intensity distribution shown as # of Mangos as approximated using background subtraction of 150 a.u. and division of the calculated mean step size of 40 a.u (**Fig. 2e**). Following normalisation the distribution was fit with three Poisson distribution fit as described in the materials and methods. (**g**) Distribution of nuclear (blue) and cytoplasmic (orange) single molecule velocities for the mCherry-M2 x24 construct, n = 229 and 224 respectively. (**h**) Distribution of nuclear (light blue) and cytoplasmic (green) single molecule velocities for the phage-CMV-CFP-MS2SLx24-tdMCP-EGFP construct, n = 461 and 163 respectively.

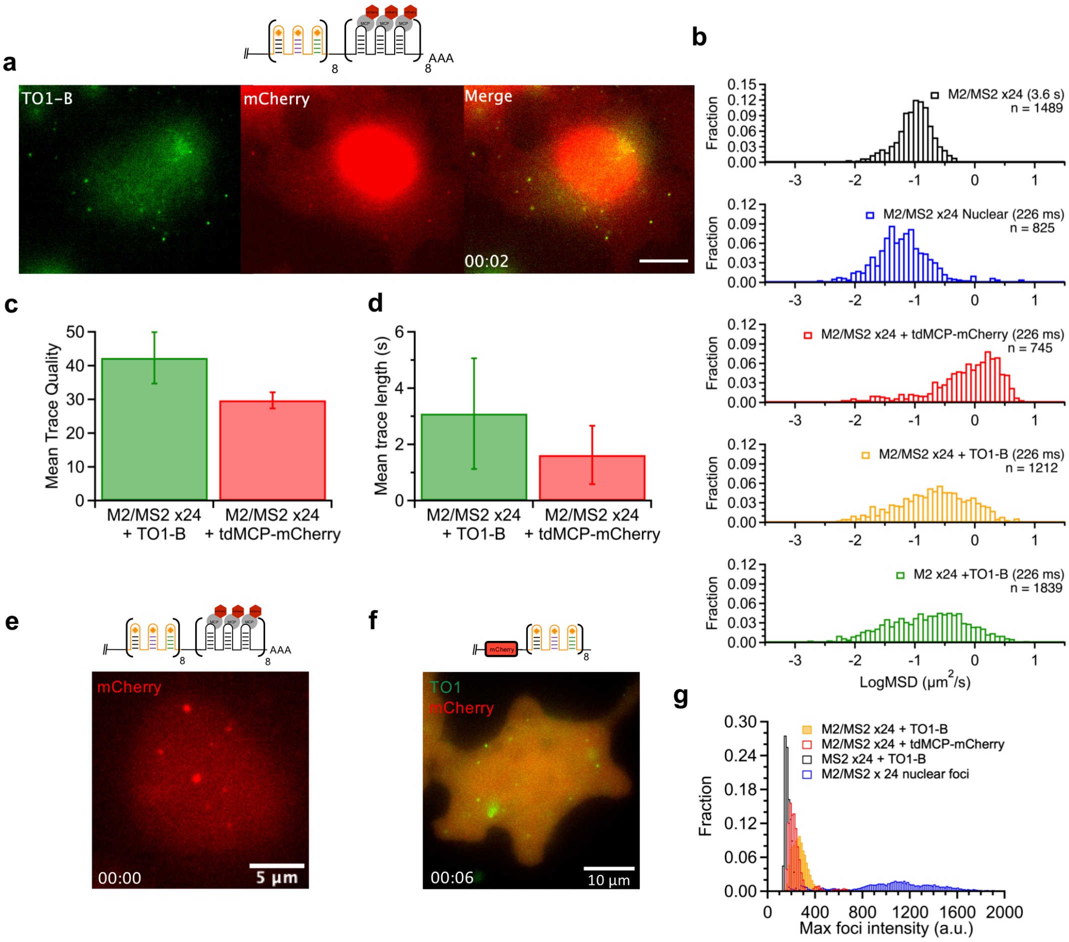

**Fig S3 | Live-cell imaging of M2 x24 and MS2-SLx24-tdMCP-mCherry**

(**a**) Live-cell snapshot of a cell expressing the M2/MS2-SLx24-tdMCP-mCherry construct depicting colocalization of the two signals. TO1-B channel – green, mCherry channel – red, scale bar = 10 µm and time shown in seconds. Still taken from supplementary video 2. (**b**) Distributions of LogMSD for live cell trajectories for M2/MS2-SLx24 tdMCP-mCherry at 3.6 s frame rate (black), nuclear M2/MS2-SLx24 tdMCP-mCherry foci at 226 ms frame rate (blue), cytosolic M2/MS2-SLx24 tdMCP-mCherry foci at 226 ms frame rate (red), M2/MS2-SLx24 + TO1-B at 226 ms frame rate (yellow) and mCherry-M2 x24 + TO1-B at 226 ms frame rate (green). N shown as the number of trajectories analysed from a total of 26 cells. (**c**) Mean trace quality as determined from the TrackMate plugin for M2/MS2-SLx24 RNA in the presence of TO1-B and tdMCP-mCherry. Green bar represents 1212 traces analysed in the TO1-B channel (green). Red bar represents 745 traces analysed in the mCherry channel (red). (**d**) Mean trace length as determined from the TrackMate plugin for M2/MS2-SLx24 RNA in the presence of TO1-B and tdMCP-mCherry. Green bar represents 1212 traces analysed in the TO1-B channel (green). Red bar represents 745 traces analysed in the mCherry channel (red). Error bars depict standard deviation of the data sets. (**e**) Representative image of M2/MS2-SLx24 + tdMCP-mCherry nuclear foci observed in the red channel, scale bar = 5 µm and time shown in seconds. Still taken from supplementary video 4a. (**f**) Representative snapshot of mCherry-M2 x24 foci in live cell showing presence of foci fluorescing specifically in the green channel and mCherry signal in the red channel, scale bar = 10 µm and time shown in seconds. Still taken from supplementary video 1a. (**g**) Live-cell foci intensity distributions for mCherry-MS2v5 x24 + TO1-B (black), M2/MS2-SLx24 + TO1-B (yellow), cytosolic M2/MS2-SLx24 + tdMCP-mCherry (red) and nuclear M2/MS2-SLx24 + tdMCP-mCherry (blue). n = 33,370, 17,544, 5881 and 12, 937 foci from a total of 26 cells. Data adapted from main figure 3 h for comparison with nuclear foci distribution.

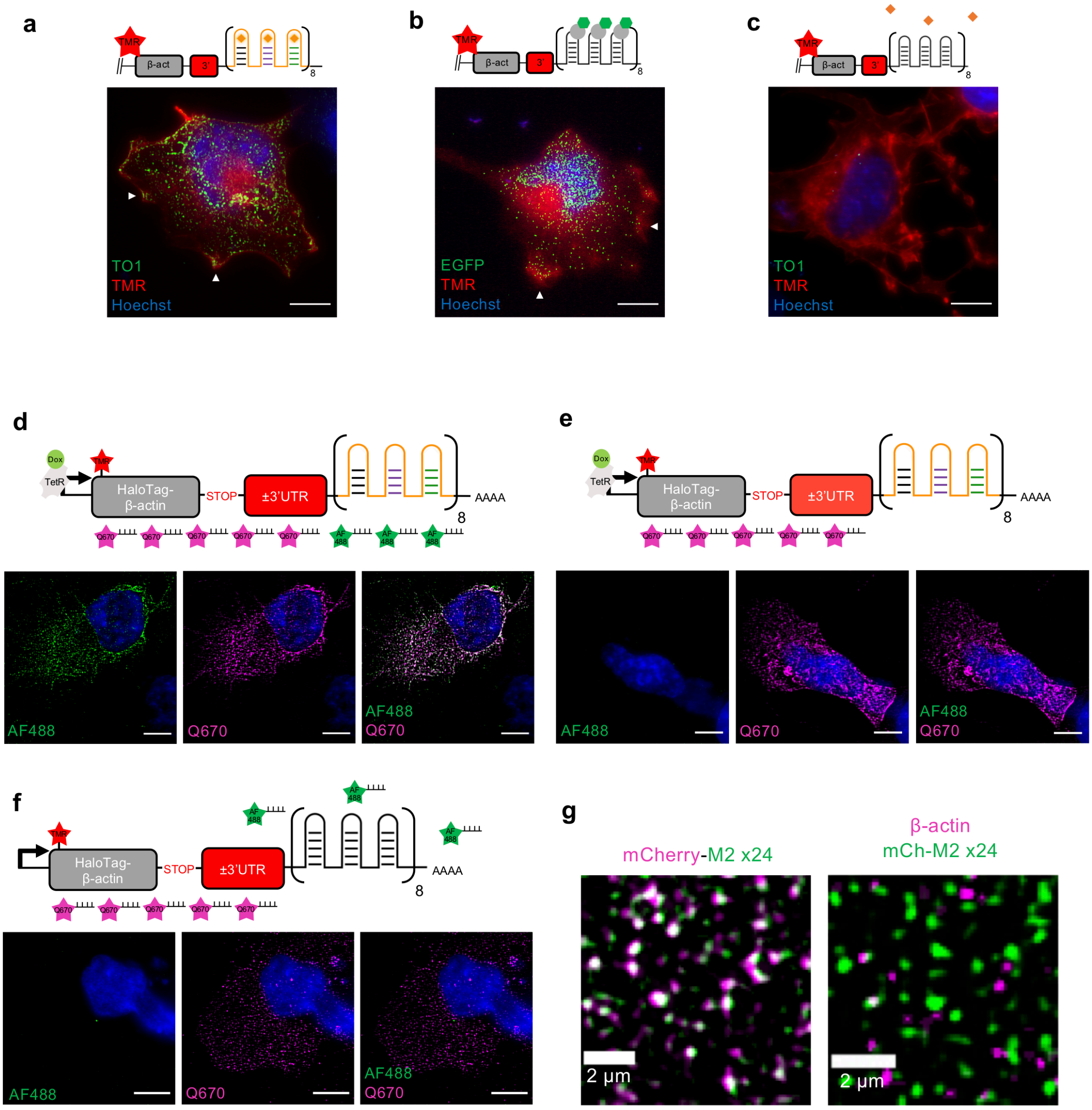


### Fig S4 | Validation of β-actin mRNA constructs

(**a - c**) Expression of Halo-tagged β-actin mRNA constructs in Cos-7 cells labelled with TMR-Halo ligand showing local expression of β-actin protein. (a) depicts Halo-β-actin-3'UTR M2 x24 + TO1-B expression, (b) Halo-β-actin-3'UTR MS2v5 x24 + tdMCP-EGFP and (c) Halo-β-actin-3'UTR MS2v5 x24 + TO1-B. White arrows depict mRNA signal at cell periphery, mRNA signal – green, protein signal – red, nuclear signal – blue and scale bars = 10 µm. (**d**) Diagram and images of Halo-β-actin-3'UTR M2 x24 expression dually labelled with β-actin-Q670 (magenta) and M2-AF488 (green) probes. (**e**) Diagram and images of Halo-β-actin-3'UTR M2 x24 expression singly labelled with β-actin-Q670 (magenta) probes. (**f**) Diagram and images of Halo-β-actin-3'UTR MS2v5 x24 expression dually labelled with β-actin-Q670 (magenta) and M2-AF488 (green) probes. Scale bars = 10 µm. (**g**) Images of mCherry-M2 x24 expression dually labelled with mCherry-Q670 (magenta) and M2-AF488 (green) probes, left and dually labelled with β-actin-Q670 (magenta) and M2-AF488 (green) probes, right. Scale bars = 2 μm.

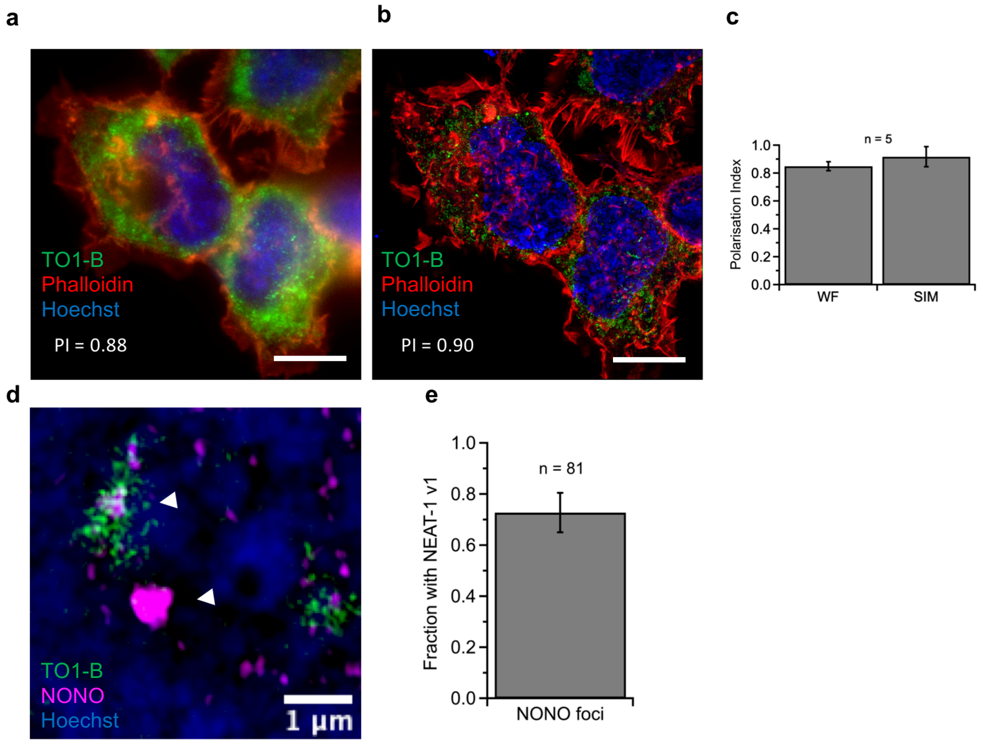


### Fig S5 | Quantification and validation of SIM images

(**a**) Widefield image of β-actin-Δ3'UTR-M2 x24 + TO1-B with average PI value depicted in the bottom left hand corner (**b**) Structured illumination microscopy image of β-actin-Δ3'UTR-M2 x24 + TO1-B cell with the average PI also depicted in the bottom left corner. Data adapted from main figure 5 for direct comparison of PI values, scale bars = 10 µm. (**c**) Quantification of the average PI value for 5 cells using both widefield and SIM images showing only a minimal change in PI. (**d**) Image of paraspeckle formation either containing NEAT-1 v1-M2 x24 (top left foci) or showing its absence (bottom left foci) as depicted with white arrows. NONO antibody stain (magenta), TO1-B (green) and Hoechst (blue). Scale bar = 1 µm. (**e**) Quantification of 81 paraspeckles showing an association of NEAT-1 v1-M2 x24 with ~ 70% of foci.

**Supplementary video 1a.**

Expression of the mCherry-M2 x24 construct in Cos-7 cells in the presence of 200 nM TO1-Biotin and imaged at 226 ms per frame in both the green (TO1-B) and red (mCherry) channels as described in the materials and methods. Time in min:sec format and scale bar = 10 μm.

**Supplementary video 1b.**

Expression of the mCherry-MS2v5 x24 control construct in Cos-7 cells in the presence of 200 nM TO1-Biotin and imaged at 226 ms per frame in both the green (TO1-B) and red (mCherry) channels as described in the materials and methods. Time in min:sec format and scale bars = 10 μm.

**Supplementary video 1c.**

Expression of the pHAGE-CMV-CFP-MS2-SLx24 construct in HEK293T cell in the presence of tdMCP-EGFP and imaged at 2 s per frame in the green (EGFP). Time in min:sec format and scale bars = 10 μm.

**Supplementary Video 2.**

Expression of the M2/MS2 SL x24 construct in Cos-7 cells in the presence of 200 nM TO1-Biotin and NLS-tdMCP-mCherry. Cells imaged at 226 ms per frame in both the green (TO1-B) and red (mCherry) channels as described in the materials and methods. Time in min:sec format and scale bar = 10 μm.

**Supplementary Video 3.**

Tracking of M2/MS2 SL x24 foci in Cos-7 cells in the presence of 200 nM TO1-Biotin and NLS-tdMCP-mCherry. Cells imaged at 226 ms per frame in both the green (TO1-B) and red (mCherry) channels as described in the materials and methods. Time in min:sec format and scale bar = 2 μm, blue trace represents track as computed using the TrackMate plugin.

**Supplementary Video 4a.**

Depiction of both nuclear and cytosolic M2/MS2 SL x24 transcripts in Cos-7 cells in the presence of 200 nM TO1-Biotin and NLS-tdMCP-mCherry. Cells imaged at 226 ms per frame in both the green (TO1-B) and red (mCherry) channels as described in the materials and methods. mCherry channel with enhanced contrast by ~4x in the middle right panel to show presence of low intensity cytosolic single molecules. These are not able to be visualised in the mCherry channel at low contrast settings (middle left panel). Time in min:sec format and scale bar = 10 μm.

**Supplementary Video 4b.**

Visualisation of bright nuclear M2/MS2 SL x24 foci in Cos-7 cells imaged using the NLS-tdMCP-mCherry signal. Cells imaged at 226 ms per frame. Time in min:sec format and scale bar = 5 µm.
